# Understanding wood decay diseases in western redcedar through development of ITS-sequencing and qPCR assays

**DOI:** 10.1101/2025.03.18.643970

**Authors:** Sydney Houston, Jun-Jun Liu, Mike Cruickshank, Arezoo Zamany, Isabel Leal, Cosmin Filipescu

## Abstract

Root and butt rot diseases cause high rates of wood decay in living stands of western redcedar (WRC; *Thuja plicata* Donn), one of the most valuable forest species in western North America. However, WRC susceptibility and the virulence of wood-decay-causing fungal pathogens are understudied, presenting a high risk for the WRC forest industry. To evaluate susceptibility of WRC to root and butt rot diseases and decay incidence, four pathogenic fungi, including *Armillaria ostoyae*, *Coniferiporia weirii*, *Heterobasidion occidentale*, and *Perenniporia subacida*, were used to inoculate WRC seedlings using two artificial methods. Next-generation sequencing (NGS) of internal transcribed spacer (ITS) region of the nuclear ribosomal DNA (rDNA) and quantitative polymerase chain reaction (qPCR) assays were developed to evaluate successful infection through detection of targeted pathogens inside the WRC tissues while development of disease was assessed by visual observation of wood decay. Disease incidence rates ranged from 20% to 60% while infection rates ranged from 80% to 100%, validating the effectiveness of the inoculation protocols. The qPCR assays designed with species-specific primers were validated for quantification of absolute abundance of *C. weirii* inside WRC host tissues with high sensitivity and specificity. These newly developed qPCR assays provide rapid, cost-effective, and accurate tools to detect early infection and latent infection in asymptomatic trees, with wide potential applications for surveillance of *C. weirii*-caused wood decay disease in greenhouse and field studies, as well as for screening of disease resistance in WRC breeding.

## Introduction

Western redcedar (WRC, *Thuja plicata* Donn ex D. Don) is an important forest species across the Pacific Northwest America with large ecological, cultural, and economic values (Antos et al. 2016). British Columbia (BC) of Canada has the world’s largest standing WRC stocks, and WRC has over $1 billion in economic activity annually in the BC forest industry (Sturrock et al. 2017; Gregory et al. 2018). Although this conifer species has shown high level of resistance to root and butt rot diseases compared to other conifers, high disease incidence still causes extensive heartwood damage in BC WRC populations. As WRC stands mature, disease incidence rates increased from ∼18% by age 50 years to 35% by age 100 of BC coastal stands, 80% by age 100 of BC interior stands, and close to 100% by age 300-400 years of both BC coastal and interior stands (Buckland 1946) as well as in Southwest Alaska (Kimmey 1956).

More recent surveys found similar disease incidence rates in BC interior sites, at ∼18% in a plantation of 48∼49-year-old trees (Cruickshank et al. 2018), and ∼75% by age 100 (Morrison et al. 2001). This is confirmed in another study that found that almost all WRC trees sampled in BC had some heartwood rot (Konchalski 2015). These increased wood-decay rates over years indicate the possibility of latent infection of decay-causing pathogens in decay-free asymptomatic trees WRC trees.

High incidence of root and butt rot diseases and extensive damage have resulted in large economic losses for the forest industry (Sturrock et al. 2017). Wood decay-caused loss accounted for up to 30% of total gross wood volume for WRC (BC Forest Service 1957; USDA 2012), which is higher than the average decay loss of other conifers, recorded at 12% (BC Forest Service 1957). In BC stands of 350 years or more, merchantable volume loss can range from 0.5 to 0.7 per stem of 1 m^3^ (Renzie and Han 2001). Although the wood decay does not kill the trees outright, it decays the lower portion of the bole, causing stem breakage by winds. However, the greater problem is economic. In addition to merchantable volume loss, decay increased harvesting costs through stem breakage after falling or skidding (Renzie and Han 2001; Sturrock et al. 2017).

Conifer root and butt rot diseases are caused by several soil fungal pathogens, commonly including *Armillaria ostoyae* (Redhead et al. 2011), *Coniferiporia weirii* (Murrill) (previously named as *Phellinus weirii* or *Poria weirii*, Zhou et al. 2016), *Heterobasidion occidentale* (Otrosina and Garbelotto 2010), and *Perenniporia subacida* (Peck). These fungal pathogens can infect living WRC trees successfully at all ages, and WRC stands have shown susceptibility to infection of these pathogens, resulting in wood decay at varying degrees (Allen et al. 1996; Sturrock et al. 2017). Most wood-decay fungi have been considered as wound parasites with their infection occurring through basal wounds and injured roots, or, through contact with infected roots (Hadfield et al. 1986). Injured roots were determined as important entry points for the decay-causing pathogens to invade, especially in young living WRC trees (Buckland 1946). A survey of interior WRC found that *A. ostoyae* was likely responsible for many of the basal scars on WRC that act as an infection court for other butt rot fungi (Cruickshank et al. 2018).

*C. weirii* was rated as causing the most serious WRC decay, and as the most common pathogen that causes wood decay in WRC living stands (Allen et al. 1996). Infection by *C. weirii* causes cedar laminated root and butt rot in WRC and yellow-cedar (*Callitropsis nootkatensis* D. Don) across western North America, as well as other species of Cupressaceae in Asia (Zhou et al. 2016; EFSA Panel on Plant Health 2018). A field survey of wood decaying fungi found that occurrence of *C. weirii* was restricted to WRC living stands with incident frequencies > 40% in Idaho (Hobbs and Partridge 1979). *C. weirii-*caused wood decay usually extended 2–3 m up the boles with the most extreme cases reaching 10 m in living WRC trees (Hagle 2006). Timber loss from the laminated root-rot disease was estimated at 4.4 million m^3^ per year in the northwestern regions of North America (Nelson 1981).

*A. ostoyae* is pathogenic to several important commercial conifers (including WRC), causing Armillaria root disease (Morrison and Pellow 2002). Conifer trees below the age of 15 years have been shown to be highly susceptible to *A. ostoyae* (Morrison et al. 1992) and Armillaria root disease killed up to 32.2% of the crop trees in young stands with an average mortality rate of 11.4% (Cleary et al. 2021). *H. occidentale* causes Annosus root and butt rot disease to a diverse group of conifers as well as numerous deciduous trees in western North America (Garbelotto and Gonthier 2013). WRC was considered as one of the living hosts (Sturrock et al. 2010), but its stumps are rarely susceptible to *H. occidentale* (Morrison et al. 1986). *Perenniporia subacida* was recently renamed as *Poriella subacida* (Gen. & Comb Nov.) based on multigene phylogenies (Chen et al. 2021). This pathogen causes white rot in both conifers and hardwoods in temperate and tropical forests. *P. subacida* was rated as one of four important white rot decay fungi on WRC in some localized areas, and BC coastal WRC was identified as its most important host (Buckland 1946).

Despite large economic loss from wood-decay fungi, little is known about infection biology and butt rot disease development of WRC trees, especially in seedlings and young trees (Hagle 2009; Cleary et al. 2012; Sturrock et al. 2017). To date, managing decay in living WRC has rarely been explored due to lack of effective tools (Sturrock et al. 2017). Developing detection tools for wood decay in living trees allows for precise estimates of incidence of disease infection, predication of mortality, accurate timber valuations, and wildlife tree assessments.

Molecular diagnostics, including quantitative polymerase chain reaction (qPCR) and microbiome profiling by next generation sequencing (NGS) of the internal transcribed spacer (ITS) region of the nuclear ribosomal DNA (rDNA), have been widely explored for monitoring the dynamics of pathogens (Stephen 2010; Schoch et al. 2012), especially at the early infection stages before disease symptoms, host defences, and impacts on tree growth are visible.

Therefore, the objectives of this investigation were to develop and assess inoculation methods for common WRC root and butt rot pathogens and detection methods using qPCR and ITS-NGS for wood decay-causing fungal species, with a focus on *C. weirii*.

## Materials and Methods

### Plant materials

Four seed families of western redcedar (WRC; *Thuja plicata* Donn) were used in this study (Supplementary Table S1). The inoculation Yr-2017 trial included two full-sib seed families #8422 (#584 x #658) and #8423 (#658 x #786), provided by the Cowichan Lake Research Station (Cowichan Lake, BC). Their one-year-old seedlings were planted in one-gallon pots using a peat-vermiculite-sand (1:1:1) mix in the spring of 2016. A 1” PVC tube was inserted offset from center and vertically into the container to accommodate the stick from the inoculation unit.

The inoculation Yr-2021 trial included two open-pollinated operational seed lots #63522 and #63738, provided by Western Forest Products (Vancouver, Canada).

### Fungal inoculation procedures

Four fungal species: *Armillaria ostoyae* (Romagn.) Herink, *Coniferiporia weirii* (Murrill) L.W. Zhou & Y.C. Dai., *Heterobasidion occidentale* (Fr.) Bref. *sensu lato*, and *Perenniporia subacida* (Peck) Donk, were used for greenhouse inoculation tests (Supplementary Table S2). Fungi of interest were cultured on 3% malt extract agar media in petri dishes. Cultures were allowed to grow until the mycelium covered most of the plate and then used as initial inoculum for the dowel or block-stick units. The fungus was easy to culture on petri plates and on autoclaved wood blocks.

Two methods, termed as the block and dowel-plug methods, were used for inoculation of seedlings (Fig. 1-A and B; Supplementary Table S1). Inoculum units for the block inoculation method were prepared following a previous protocol (Sturrock and Reynolds 1998; Cruickshank et al. 2020) with modification. In brief for the blocks used in the block-stick or block without stick units, stems of juvenile red alder (*Alnus rubra* Bong.) were harvested and cut into wood block segments with ∼ 5 cm diameter x ∼ 6 cm length. A hole of 9.5 mm diameter was drilled longitudinally 3-4 cm through the centre of each stem segment. Twenty to 25 wood blocks were then placed side by side into an autoclavable bin. Distilled water was added to each bin so that the top of the wood blocks was just covered with water. The bin was then autoclaved for 75 min at 121°C. After autoclaving, excess water was drained, and 8 wood blocks were placed in two stacks of 4 with the drilled hole side pointed up. Both stacks of wood blocks were placed in a 20 x 56 cm autoclavable mushroom spawn bag with 0.2-micron microporous patch to allow gas exchange. The tops of bags were left open and then autoclaved for 75 minutes at 121°C slow exhaust cycle. After autoclaving, the tops of bags were folded over and held with 1 inch fold back clips, and then cooled in a sterile flow hood overnight. Wood bags were inoculated aseptically by placing 2 cm square cubes from half of a fungal colonized petri plate per bag, distributed evenly on the edge of each wood block. The inoculated bags were re-clipped and placed in the dark at 10°C in a walk-in cooler to allow the fungus to grow for up to one year.

**Figure 1.**
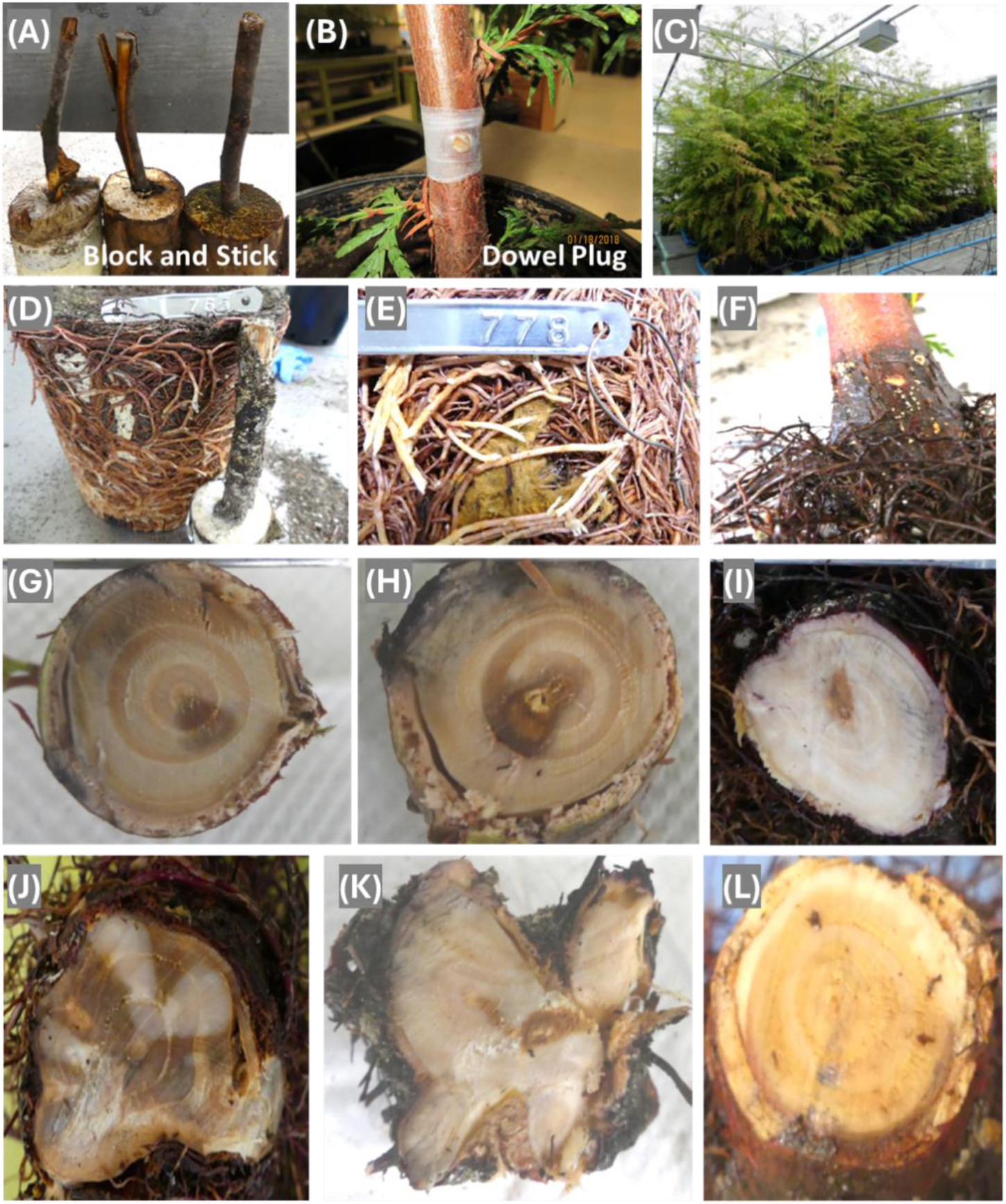
Diseased symptoms of western redcedar caused by root- and butt-rot fungi. Assessment was performed at 18-months post controlled inoculation. (A) Inoculums prepared using the block and stick method, they were inserted into pots for inoculation of roots; (B) Inoculation of *Armillaria ostoyae* using the dowel-plug method; (C) Growth of seedlings post inoculation under controlled conditions in a greenhouse; (D-E) Mycelia on the fine root surface of seedlings inoculated by *Perenniporia subacida* and *Coniferiporia weirii*, respectively; (F) Mycelium of *P. subacida* on coarse roots and root collar ; (G-H) Wood decay symptoms caused by *C. weirii* using the dowel-plug method; (I-L) Wood decay symptoms of western redcedar root collars of seedlings inoculated with *P. subacida*, *A. ostoyae*, *C. weirii*, and *H. occidentale*, respectively, using the block-stick method.

The block-stick transfer sticks were prepared using two-year-old branches of various species (western red cedar, western hemlock, and Garry oak). These branches were harvested and cut into sticks with diameter at one end slightly larger than the block hole. They were rinsed lightly with tap water and allowed to air dry. Once the alder wood blocks were colonised, a WRC living stick was inserted into each hole of the blocks inoculated with *C. weirii* and *P. subacida*; living Garry oak sticks were used for blocks inoculated with *A. ostoyae*; living hemlock branch sticks were used for blocks inoculated with *H. occidentale*. Sticks were inserted by trimming one end, so it fit tightly into the block hole. The branch sticks were trimmed 1” shorter than the depth of the pot. The block end was covered with a plastic bag, taped closed, and aluminum foil was placed over the block to keep it dark. After removal of the PVC tube from the seedling pots, the stick of the unit was inserted into the hole left behind by the PVC tube in each pot, with the block remaining above ground. Excess space around the branch stick was filled with soil.

For the dowel-plug inoculation method, inoculum units were prepared using wood dowels of stem cross sections of 8 mm, and they were obtained from a 35-year-old western red cedar tree in the fall of 2017. Dowel plugs at 5 mm diameter x 3 mm thickness were prepared from the heartwood using an increment borer. The dowels were autoclaved in bags for 30 min, cooled in a laminar flow hood, and placed on colonized petri dishes until the wood dowels were colonized. An increment borer was used to remove a bark/stem section from each of the greenhouse trees between internodes at approximately 10 cm from the soil. A colonized dowel- plug was inserted and trimmed to fit flush with outer bark. The inoculation point was wrapped with parafilm around the stem and tied with grafting rubber to hold the plug-in place. For controls of non-inoculated trees, an un-colonized plug was inserted.

For the inoculation Yr-2017 inoculation trial (Supplementary Table S1), two-year-old seedlings of families #8422 and #8423 were moved into two-gallon pots in the spring of 2017 and inoculated in November 2017 (year 3) using either block-stick or dowel-plug method.

Disease symptoms were assessed, and WRC tissues were sampled at 18 months post inoculation in June 2019. For the inoculation Yr-2021 trial (Supplementary Table S1) to detect early infection by *C. weirii*, two-year-old seedlings of two open-pollinated seedlots #63522 and #63738 were inoculated in Dec. 2021 using a block-without-stick method. These one-year-old nursery seedlings were grown in one-gallon plots for one year in the same soil mix. In the fall of the second year, the pots received a colonised block without a stick that was kept in the cooler from the 2017 experiment. The block was placed with the block end (flat surface) on the top of the soil and a plastic bag was taped around the block to prevent desiccation. Soil was added as necessary to make contact with the circumference of the block so there were no air spaces. The flat end touching the soil was not covered with the bag to allow the fungus in the block to have direct contact with the soil. The block was then wrapped with aluminum foil to keep it dark, and then masking tape was placed over the top end of the block, completely around the pot underside a few times to hold the block firmly in place on the soil surface. Care was taken not to disturb the block. Disease symptoms were assessed, and fine roots were individually sampled from each seedling at six-months post inoculation in July 2022.

In all cases the soil in the pots was kept moist but not wet. In many cases, the fungus could be seen moving across the top of the soil from the block and sometimes a short way up the stem of the tree. The fungal movement on the soil occurred in the fall when the days became shorter. It was important to keep the soil surface from completely drying out at this time. The trees in the Yr-2017 trial were placed on a bench in a temperature-controlled glass greenhouse kept between 12-18°C. The temperature was controlled by forced air and evaporative coolers. The trees in the Yr-2021 trial were placed in a plastic shade house on a bench with summer temperature controlled by fan and overhead and side vents ranging from 2-35°C. The shade house and glasshouse were heated in winter, so the temperature never went below 0°C. Trees in the Yr-2017 trial were automatically watered with drippers and the trees in Yr-2021 trial were watered by hand. No additional fertilizer was added, and the trees had no other pests.

### Symptom assessment of fungal infection

At the end of the inoculation trials, seedling heights and stem diameters at ground level were measured. Stem (including bark) diameter was measured for each seedling at the soil line using a Vernier caliper at two positions 90° apart. Inoculum units used for the block-stick method were checked visually for fungal presence in the wood blocks and sticks. Inoculum units were considered as successful if ectotrophic mycelium had grown at least half-way down along the sticks of branch segments and transfer of fungal pathogens to the seedlings was confirmed by presence of ectotrophic mycelium on roots (Sturrock and Reynolds 1998). The mycelium on the roots and stick was distinct and easily seen visually. For the block without stick, only wood blocks were checked for fungal growth.

For destructive evaluation of fungal infection, stems of seedlings inoculated by the dowel-plug method were cross-sectioned at the inoculation sites. For the seedlings inoculated by the block or block-stick method, sampling was done at the root collar of the soil lines. The root collar is the lower part of the stem at the soil line where the roots arise. Stem cross-sections at the soil line or in large roots near the collar were checked for the presence or absence of dark staining and decay inside the wood tissues as an indicator of xylem decay caused by root rot pathogens (Sturrock and Reynolds 1998).

To detect decay fungi in WRC tissues, wood (including bark) samples were taken from all inoculated and control seedlings at the root collar for both block-stick methods or the stem at the wound-inoculation area for the dowel-plug method. Fine roots were individually collected from those seedlings inoculated using the block-stick method.

### gDNA extraction

A DNeasy Plant Pro Kit (Qiagen) was used for extraction of gDNA from fine roots and fungal mycelium, following the manufacturer’s protocol with modifications: an initial step of homogenizing freeze-dried tissue was performed using a Fisherbrand™ Bead Mill 24 Homogenizer (Fisherbrand) under liquid nitrogen using the conditions of speed = 5 m/sec, time = 7 sec, cycle = 1, and dwell = 0, and repeating this step if necessary. A 15 min incubation at 65°C with shaking at 5 min intervals was added to solubilize the cell wall and release intracellular components. Washing steps were repeated twice to remove the lysis reagents. The elution buffer was warmed to 65°C and kept on the column for 5 min prior to elution.

A hexadecyltrimethylammonium bromide (CTAB)-based protocol (Healey et al. 2014) was followed with modification to extract DNA from stem samples with wood and bark tissues. In brief, stem samples were freeze-dried, ground into fine powder using liquid nitrogen, and mixed with preheated extraction buffer (2% CTAB; 200 mM Tris, pH 8.0; 50 mM EDTA, pH 8.0; 1 M NaCl; 0.5% activated charcoal; 1.5% polyvinylpolypyrrolidone; and 1.5% β-mercaptoethanol). The mixture was incubated at 65°C for 30 min, and then extracted twice with an equal volume of chloroform. Following centrifugation, the upper phase was recovered and mixed with an equal volume of 100% isopropanol to precipitate nucleic acids. After centrifugation, the pellet was re-suspended in a DNA resuspension buffer (1 M NaCl; 0.5% SDS; 10 mM Tris-HCl, pH 8.0; 1 mM EDTA, pH 8.0; and 10 µg/mL RNase A) and incubated at 65°C for 10 min. The resuspension solution was extracted using phenol: chloroform: isoamyl alcohol (25:24:1, v/v), and followed by extraction with chloroform. DNA was precipitated by mixing with two volume of 95% ethanol. After centrifugation, the pellet was washed in 70% cold ethanol, dried, and re- suspended in nuclease-free water.

The integrity of the extracted DNAs was checked using agarose gel electrophoresis. DNA purity and concentration were measured using a NanoDrop™ 2000/2000c Spectrophotometer (Thermo Fisher Scientific) and a Qubit dsDNA Broad Range Assay Kit on the Qubit Fluorometer 3.0 (Thermo Fisher Scientific).

### NGS analysis of the rDNA-ITS regions

The abundance of four fungal species were measured in cedar tissues through fungal ITS-NGS using Illumina MiSeq platform. To avoid amplification of plant gDNAs, a nested PCR approach was used to amplify fungal ITS1 sequences (Adamo et al. 2020). For the 1st round PCR, reaction mixture was prepared in a 20 µl volume per reaction with Premix Taq™ DNA Polymerase (Ex Taq™ Version 2.0) (Takara Bio Inc., San Jose, USA) master mix, fungal-specific primers ITS1F and ITS4 (White et al., 1990; Gardes and Bruns 1993; Supplementary Table S3), and 20 ng of DNA extracted from cedar tissues. The 1st round PCR was run with initial denaturation at 95°C x 3 min; 30 cycles of 95°C x 30 sec, 50°C x 30 sec, 72°C x 60 sec; followed by a final extension at 72°C x 7 min.

After completion of the 1st round of PCR, the PCR products were diluted 10 X and used for the 2nd round PCR to prepare ITS-amplicon libraries using primers ITS5 and ITS2 that targeted fungal ITS-1 region (White et al., 1990; Supplementary Table S3). For preparation of the ITS-amplicon library, ITS5 and ITS2 were tagged with primers TruSeqF and TruSeqR (Illumina), respectively. For the 2nd round PCR, reaction mixture was prepared with 10 X PCR buffer with 15mM MgCl2 (Qiagen), DMSO (Roche), dNTP mix 10 mM (New England Biolabs), FastStart High Fi DNA polymerase (5U/µl, Roche). The 2^nd^ PCR run was with an initial DNA denaturation at 96°C x 15 min followed by 35 cycles of 96°C x 30 sec, 52°C x 30 sec, and 72°C x 60 sec; with final extension at 72°C x 10 min. The fungal ITS amplicons were verified by electrophoresis on 1% agarose gels. After adding barcoding oligonucleotides using Dual-Indexes (Integrated DNA Technologies) and Illumina adapters to each sample, the amplicon was quantified using a Quant-iT™ PicoGreen® dsDNA Assay Kit (Life Technologies) and multiple samples were pooled in equimolar concentration as the ITS-amplicon library. After cleaning up using sparQ PureMag Beads (Quantabio), the ITS-amplicon library was sequenced using a MiSeq Reagent Kit (v2, 500 cycles) on the Illumina MiSeq platform at Génome Québec Innovation Centre, Canada.

### Quantitation of the targeted fungi using ITS-NGS reads

FASTQ files of 2 x 250-bp paired-end (PE) reads were imported into CLC Genomics Workbench (ver 23.0, Qiagen) with removal of failed reads and retaining quality scores (NCBI/Sanger or Illumina 1.8 pipeline or later). The Operational Taxonomic Unit (OTU) Clustering (ver 2.6) in the CLC Microbial Genomics Module (ver 23.0) was used for taxonomic classification of fungal microbiome. Clean reads of each sample were classified into OTUs at 97% sequence similarity using UNITE-reference database (https://unite.ut.ee/#main; Sh_general_release_seq161,355; 16.10.2022). OUT relative abundance at threshold of 0.01% (target reads in total ITS reads) was considered for presence of pathogens for their successful infection. The relative abundance of the targeted fungi was calculated as percentages of the targeted fungal reads in all clustered reads for each cedar tissue sample.

### Development of species-specific qPCR assays

*C. weirii* was selected to develop qPCR assays because this fungus was the main focus of our screening of WRC genetic resistance against the root and butt rot diseases. Six nuclear gene sequences were selected for design of *C. weirii* species-specific primers, including ITS, small and large subunits of ribosomal RNA (MS and ML), the largest and the 2nd largest subunits of RNA polymerase II (RPB1 and RPB2), and translation elongation factor 1-alpha (Tef1). These DNA sequences were retrieved from the NCBI GenBank (http://www.ncbi.nlm.nih.gov/genbank) and aligned with related sequences of other fungal species using Clustal Omega (https://www.ebi.ac.uk/jdispatcher/msa/clustalo). *C. weirii* DNA regions highly variable across different fungi were targeted to design PCR primers using ThermoFisher Primer Express Software 3.0.1 (Supplementary Table S3). All primer pairs were newly designed in this study except for Cw-ITS primers (F160/R683), which were modified from a previous report (Lim et al. 2005), adding additional nucleotides at the 5’-end of the original sequences to increase primer annealing temperature to 60°C for improved resolution based on the *C. weirii* genome sequence available recently (McMurtrey et al. 2023).

Specificity of *C. weirii* species-specific primers was first tested using endpoint PCR using gDNA extracted from cultured mycelia of five decay fungi, including *C. sulphurascens* (Supplementary Table S2). Endpoint PCR reaction solution contained 5µl Premix Taq™ DNA Polymerase (Ex Taq™ Version 2.0) (Takara Bio Inc., San Jose, USA) master mix, 1µl of each PCR primers (10 mM), and 1 ng of gDNA from each individual fungus for a total volume of 10 µl per reaction. Endpoint PCR was run using a Veriti^TM^ 96-Well Fast Thermal Cycler (Applied Biosystems, Thermo Fisher Scientific), with conditions programmed as an initial DNA denaturation at 95°C for 10 min, 40 cycles of 95°C x 40 sec, 58°C x 20 sec, 72°C x 30 sec; followed by a final extension at 72°C x 9min. Amplicons of endpoint PCR were checked by electrophoresis in a 1% agarose gel containing 0.05% RedSafe (FroggaBio). DNA fragments were visualized and photographed under ultraviolet light using a gel documenter (Syngene, USA). The amplicon sizes were estimated by comparison with DNA standards. In addition, qPCR was also performed to check species-specificity of the PCR primers using 1 ng of individual fungal gDNA with conditions as described below.

Efficiency of *C. weirii* species-specific qPCR assays were assessed by analysis of qPCR standard curves. *C. weirii* pure gDNA samples were prepared by performing 10 X serial dilutions to obtain DNA concentrations in a range of 20 ng/µl to 0.2 pg/µl. Each qPCR mixture was set up using a 0.2 mL 96-well plate, containing 10 µL 2 X SYBR™ Select Master Mix (Applied Biosystems), 1 µl standard DNA at a corresponding concentration from the gDNA set of 10 X serial dilutions, and 1 µL of each primer (10 mM) for a total volume of 20 µl per reaction. All DNA samples were tested in triplicate with negative controls (no DNA) for at least three independent runs. The qPCR was run on a QuantStudio™ 3 Real-Time PCR System (Applied Biosystems) using standard cycling mode following the product manual. PCR conditions were set up with UDG activation at 50°C x 2 min and 95°C x 2 min, and 40 two-steps cycles with 95°C x 15 sec and 60°C x 1 min. The standard curve was generated by plotting the linear relationship between the cycle threshold (Ct) values and absolute *C. weirii* DNA amounts in each PCR reaction. The efficiency (E) of qPCR was calculated as the fraction of target molecules regenerated during each PCR cycle using the slope of the standard curve (Lalam 2006; Alvarez 2007); E = 10^−(1/slope)^ – 1. A slope value of the standard curve at –3.32 indicates a qPCR efficiency at 100%.

These qPCR assays were used to detect and precisely quantify *C. weirii* levels in WRC tissue samples collected from both infected symptomatic (decayed) and asymptomatic (decay-free) seedlings. This set of WRC tissue samples was also included in the ITS-NGS analysis described above for comparison between these two molecular diagnostic approaches. Each tissue sample was repeated in a qPCR run and their mean Ct value was used for quantitation. Based on Ct values of the tested cedar tissue samples, the absolute amount of *C. weirii* DNA was calculated for each sample using the equation of the standard curve. The relative abundance of *C. weirii* inside cedar tissues was transformed as the targeted fungal amount per total amount of tissue DNAs in pg or ng of fungal target DNA vs. ng of total DNA extracted from WRC tissues.

### Statistical analysis

Fungal infection rates were calculated based on decay development and presence of the decay fungus in WRC tissues as detected by ITS-NGS and qPCR assays. A t-test was used to test tree growth difference between controls and inoculated treatments. One-way ANOVA was used to test the difference in tree growth among tree groups based on wood-decay symptoms or presence/absence of pathogens as detected by ITS-NGS and qPCR. The multiple comparisons were corrected by the Bonferroni-Holm method.

The relative DNA abundance value of targeted fungi was calculated as the DNA amount of targeted fungus in relation to the total amount of DNA used in a qPCR assay, or the number of NGS reads of the targeted fungus in relation to the total number of NGS reads in a WRC tissue sample as detected by the ITS-NGS. The Kruskal-Wallis test was used to compare relative abundance of pathogens as measured by ITS-NGS, and multiple comparison by the Benjamini-Hochberg FDR method. To compare the effectiveness and consistency of molecular detection by ITS-NGS and qPCR, Pearson’s correlation (R value) was analyzed for the relative fungal DNA abundance values as detected by ITS-NGS and qPCR. Before Pearson’s correlation analysis, the relative fungal values were transformed using - log10 (values + 10^-6^).

## Results

### Assessment of wood decay diseases caused by four pathogenic fungi

No morphological change (such as needle color, or loss of foliage) was observed in the inoculated seedlings as compared to controls at 18-months post inoculation (Fig. 1-C), but fungal mycelia were observed on roots of some seedlings inoculated by the block-stick method (Fig 1-D to F). Wood decay was the most visual disease symptom in the inoculation trials of this study. Wood decay was not observed in any of the non-inoculated control seedlings up to 18- months post inoculation. In contrast, wood decay was observed as discoloration of tissue at the root-collar and in the wounded stem at the inoculation sites at 18 months post inoculation caused by all four pathogens (Fig. 1-G to L). In combination of data from the two inoculation methods, incidence rates of wood-decay disease in the Yr-2017 trial were determined to be 20%, 40%, 60% and 60% at 18-months post inoculation by *A. ostoyae*, *C. weirii*, *H. occidentale*, and *P. subacida*; respectively (Supplementary Table S4 and Fig. S1). By combining data from the two inoculation methods and four inoculated fungi, the full-sib families #8422 and #8423 had a total disease incidence rate of 35% and 55%, respectively, but without significant difference between them (X^2^ test, p = 0.0608).

Inoculation using the block-stick method produced decay rates ranging from 17% to 50% and the dowel-plug method produced decay rates of 0% for *A. ostoyae* and 75% for the other fungi in Yr-2017 trial (Supplementary Fig S1). Inoculation by *C. weirii* in the Yr-2021 trial produced no decay development at 6 months post inoculation using the block without stick method.

### Fungal infection detected by ITS-NGS

ITS-NGS generated a total of 18 millions of 2 x 250-bp PE reads, averaging 195K ± 33K reads with a range from 117K to 288K reads across all samples. The clean reads were annotated to 328 OTUs of the UNITE-reference database at 97% sequence similarity, including the four targeted fungal species, but *H. parviporum* (GenBank acc. X70021) was selected as the closest species because *H. occidentale* was not included in the UNITE-reference database.

At a threshold of 0.01% for relative OUT-abundance, ITS-NGS detected pathogen-positive rates ranged from 60% (*C. weirii*) to 80% (*A. ostoyae*) across all four decay fungi in all inoculated seedlings (subtotal included both decayed and decay-free seedlings) at 18-months post inoculation in trial 2017 (Fig. 2-A). In decayed seedlings, pathogen-positive rates varied from 50% (*C. weirii*) to 100% (*A. ostoyae*). The targeted fungus was below the threshold of relative abundance in tissue samples of four decayed seedlings: two infected by *C. weirii*, one infected by *H. occidentale*, and one infected by *P. subacida*, indicating that ITS-NGS did not detect the presence of pathogens in some symptomatic trees. In inoculated but decay-free seedlings, ITS-NGS detected latent infection with incidence rates in a range from 50% (*H. occidentale*) to 75% (*A. ostoyae*) at 18 months post inoculation.

**Figure 2.**
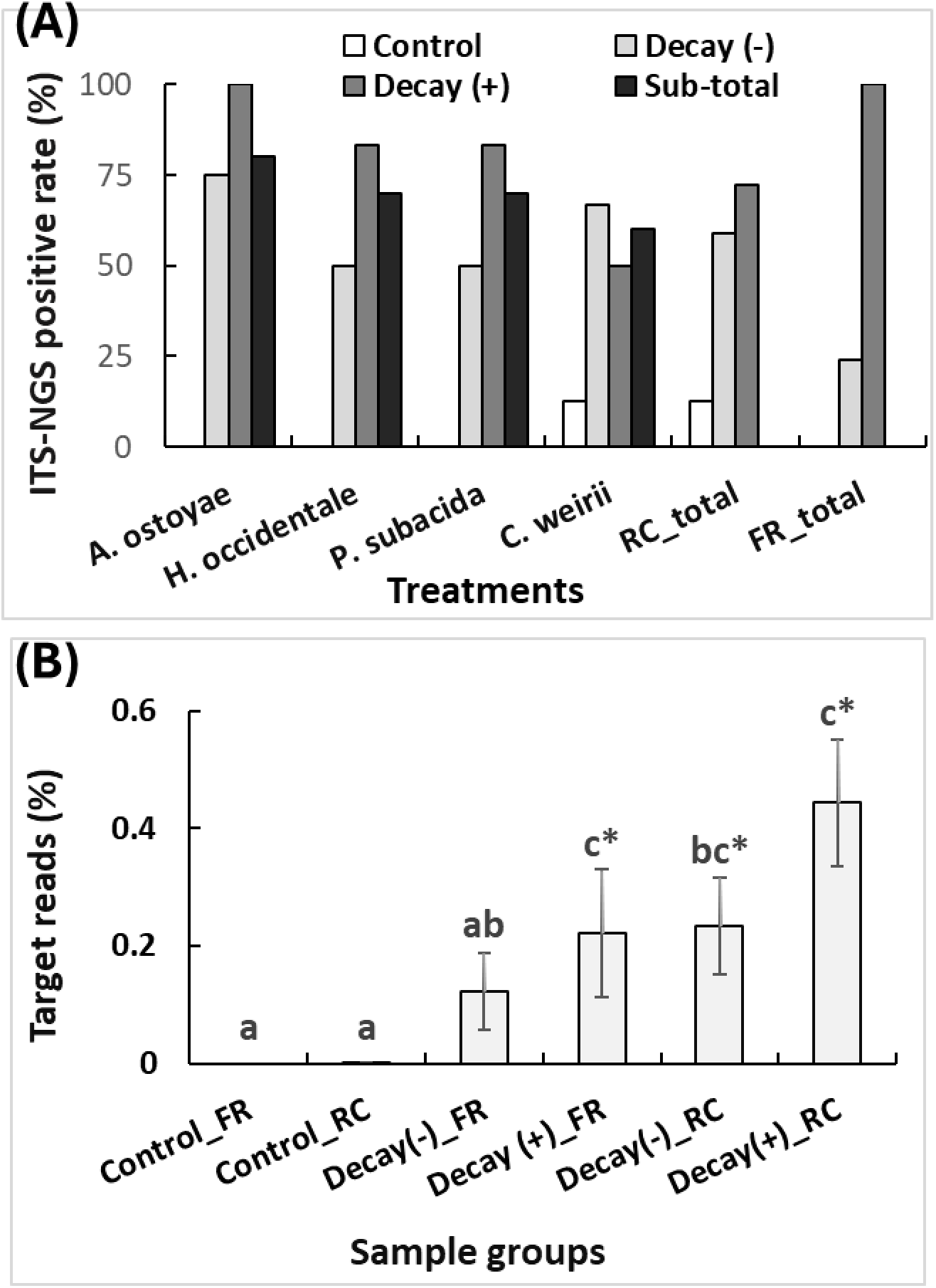
Detection of targeted decay fungi by ITS-NGS in western redcedar tissues collected at 18 months post inoculation in trial Yr-2017. Relative levels of targeted fungal reads were normalized as percentage of total reads, with a threshold at 1 x 10^-4^. Fungal infection by both block and dowel-plug methods was determined by presence of fungal sequences in the tissues of inoculated western redcedar seedlings. (A) Successful infection rates of four individual decay fungi; Decay (-): inoculated but decay-free seedlings; Decay (+): inoculated and decayed seedlings; FR are fine roots and RC is root collar. (B) Comparison of relative abundance of targeted decay fungi in western redcedar tissues. FR and RC as above; Control non-inoculated seedlings; Decay (-) and Decay (+) as above. Columns represent means of relative abundance of targeted decay fungi with standard deviation of mean (SEM). Different letters indicate significance among six groups of tissue samples as analyzed using Kruskal Wallis with multiple comparisons corrected by Benjamini-Hochberg where one star indicated p < 0.05.

ITS-NGS detected pathogen-positive rates were further compared between two WRC tissue types. In all decayed seedlings inoculated by the four pathogens, 100% of their fine root samples and 72% of their root-collar were positive for the targeted fungus (Fig. 2A; Supplementary Table S4; Fig. S2-A and S2-B). In all decay-free seedlings inoculated by the four pathogens, 59% of their stem wood and 24% of fine root samples were pathogen-positive, indicating that these decay-free seedlings had latent infection by the targeted decay fungi.

Although no decay was observed in all seedlings of the Yr-2021 inoculation trial, ITS-NGS also detected early infection by *C. weirii* in one of 10 seedlings (10% of total) at six-months post inoculation (Supplementary Fig S3-A). In contrast, none of all four targeted fungi was detected in either the fine roots or the root-collar samples of all non-inoculated seedlings with the exception of *C. weirii* being detected in the root-collar tissue of one non-inoculated seedling (Fig. 2-A; Supplementary Fig. S2-B and S3-B).

Relative ITS-NGS reads of the targeted fungi ranged from 0.014% to 99.89% of the total ITS-NGS reads across all pathogen-positive WRC tissue samples (Supplementary Fig. S2-A and B). Targeted fungal NGS reads accounted for 44% and 23% of total reads in averages of root-collar samples of decayed and decay-free seedlings, respectively (Fig. 2-B. As compared to the root-collar, fine roots had lower relative abundance levels of the pathogens, accounting for averages of 22% and 12% of total NGS reads in decayed and decay-free seedlings, respectively (Fig. 2-B.

Across all trees, the normalized fungal NGS reads showed significant correlation between fine roots and root collars of tested seedlings (Pearson R = 0.538, n=32, *p* = 0.0016, Supplementary Fig. S2C). Across six groups of tissue samples (Fig. 2 B), three groups (root collars of either decayed or decay-free seedlings, and fine roots of decayed seedlings) showed pathogen abundance significantly higher than non-inoculated controls. Pathogen abundance in fine roots of inoculated but decay-free seedlings (Kruskal-Wallis rank sum test, p < 0.05) was significantly lower than fine roots of decayed seedlings (Fig. 2-B). In summary, these analyses indicated that the detection of fungal infection by ITS-NGS was more accurate in root collar tissues than in fine roots in either decayed or decay-free seedlings.

### Development of qPCR assays for detection of *C. weirii*

Six qPCR assays (Supplementary Table S3) were designed and tested using endpoint PCR followed by agarose gel electrophoresis. It was found that four of the assays: Cw-ITS (F160/R683), Cw-ML (F88/R235), Cw-RPBa (F275/R459), and Cw-RPBb (F76/R235), were *C. weirii* species-specific, with expected amplicons exclusive in *C. weirii* and no amplification in the other fungi (*A. ostoyae, C. sulphurascens, H. occidentale*, and *P. subacida*). In contrast, the other two PCR assays Pw-MS (F234/R399) and Pw-EF (F97/R292) showed weak amplicons in other fungal species (Supplementary Fig. S4; Table S5).

These qPCR assays were further tested for their species-specificity by using 1 ng gDNA of different fungi as templates in qPCR runs (Supplementary Table S6). Three qPCR assays, Cw- RPBa (F275/R459), Cw-RPBb (F76/R235), and Cw-ITS (F160/R683), were shown to solely amplify *C. weirii* with the other fungi having negative results. These assays showed amplicon melting temperatures (Tm) (83.06°C-84.01°C) and qPCR Ct values (19.04-22.26) as expected for *C. weirii*. The Cw-MS primer pairs (F234/R399) amplified similar fragments in both *C. weirii* and *C. sulphurascens* (melting Tm = 75.06°C and 75.14°C, respectively), but Ct values (e.g. *C. weirii* at Ct = 16.59 vs. *C. sulphurascens* Ct = 36.99) were more delayed in *C. sulphurascens*. The remaining two qPCR primer pairs Cw-EF (F97/R292) and Cw-ML (F88/R235) had amplicons in *C. weirii* and other decay fungi, but *C. weirii* had much earlier Ct values (22.24 vs. 34.46-35.96; 18.93 vs. 34.23) than other fungi, and amplicon melting temperatures were different from others (86.26°C vs.80.52-83.12°C; 80.25°C vs. 85.44°C). Ct values for detection of other fungi (Ct > 34.23) were close to or outside the reliable maximum, demonstrating that these six qPCR assays were well designed for *C. weirii* species-specificity.

Based on the above assessment of qPCR assay specificity, five qPCR assays were used to generate standard curves using a set of 10 X serially diluted *C. weirii* gDNA amounts. The PCR assay Cw-RPBb (F76/R235) showed the best qPCR efficiency, resolution, and repeatability. The amplicons of this assay were linear over five levels of DNA amounts from 2 ng (100 pg/µl) to 0.2 pg (10 fg/µl) per reaction (Fig. 3-A). All qPCR reactions were highly repeatable with high amplification efficiencies (E = 0.941 ± 0.013). At the lowest concentration of 0.2 pg per reaction (20 fg/ul), *C. weirii* gDNA was reliably detected in all independent experiments (R^2^ = 0.9806) (Fig. 3-B). The qPCR performance indicated that this qPCR assay Cw-RPBb (F76/R235) was highly effective in the detection of *C. weirii*, without inhibitory effects on amplification. The other four qPCR assays had qPCR efficiency at lower levels (0.554 ≤ E ≤ 0.838) (Supplementary Fig. S5).

**Figure 3.**
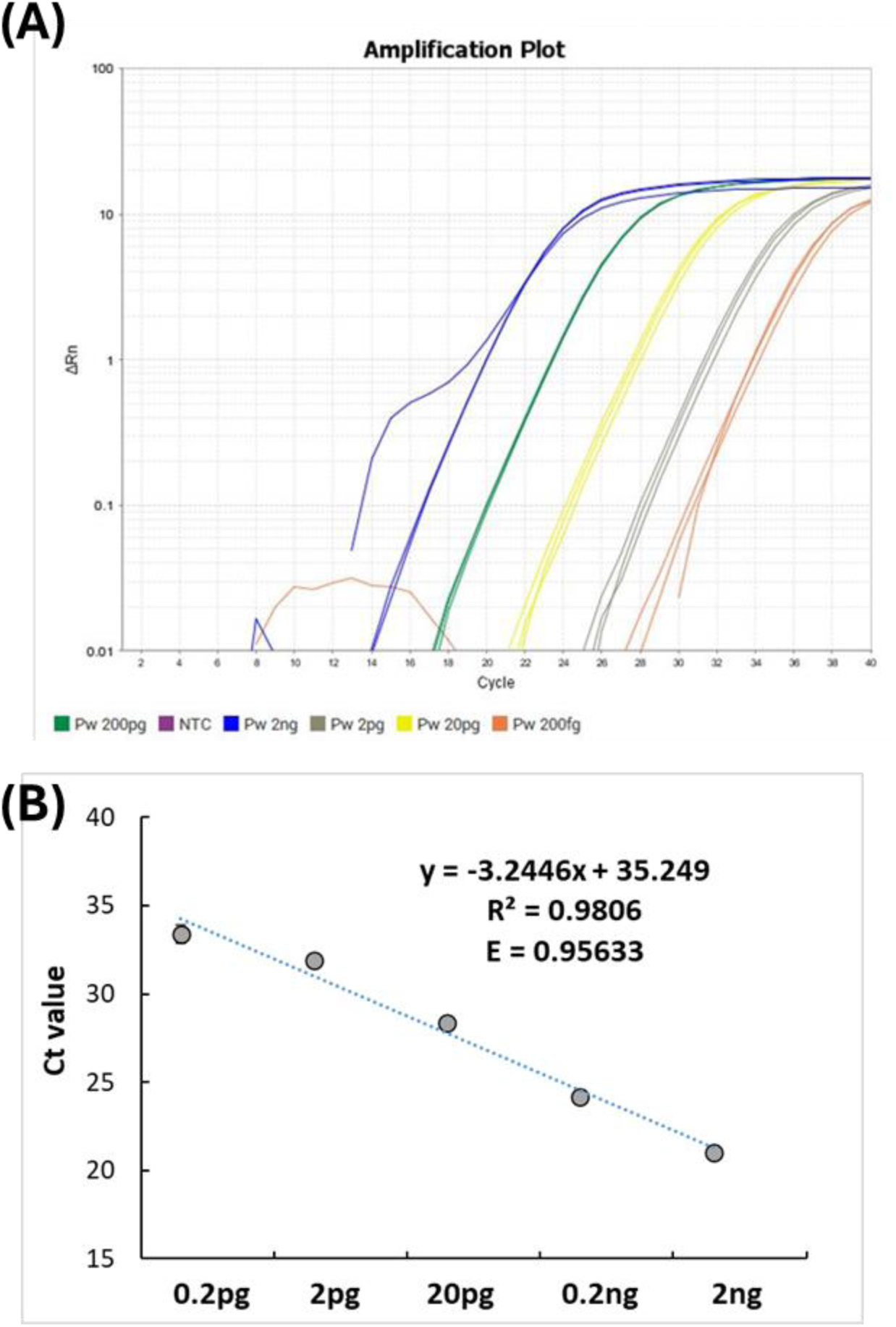
Standard curve analysis to show sensitivity, repeatability, and efficiency of qPCR assays for detection of gDNA of *C. weirii*. The qPCR assay was run using SYBR Green Master Mix with the taxon-specific primer. Each DNA dilution was tested in triplicate. (A) Amplification plot showing Ct values of pure *C. weirii* gDNA with tenfold dilutions ranging from 2 ng to 0.2 pg per PCR reaction. Absolute DNA amounts at five final concentrations (100 pg/µl to 10 fg/µl) with three repeats were tested by the qPCR assay. (B) A standard curve generated by plotting the mean Ct values versus the absolute amounts of *C. weirii* gDNA. Standard curve displaying a relationship between the log 10 of known amount (pg) of target gDNA and Ct value of qPCR amplicon. A linear relationship between log-transformed DNA amounts in PCR reactions and cycle threshold (Ct) values were detected, with qPCR efficiency E = 0.956.

### Validation of qPCR assays in inoculation trials

The qPCR assay Cw-RPBb (F76/R235) was selected to evaluate infection levels of WRC root and root collar samples. Fine roots were tested in a total of 16 seedlings inoculated by block-stick or block method. All ten seedlings from the Yr-2021 trial were visually decay-free at six-months post inoculation, but qPCR detected 30% as *C. weirii*-positive during the early infection process before symptom development. This rate of early infection detected by the qPCR assay was three times higher than that detected by ITS-NGS. In the six fine root samples collected at 18 months post inoculation from the Yr-2017 trial, qPCR showed that 33% of trees tested positive for the presence of *C. weirii*, which was higher than that (16%) detected by ITS- NGS (Supplementary Fig. S3-A). After transformation of qPCR Ct values into relative fungal gDNA amounts in proportion to total DNA extracted from fine root tissues, *C. weirii* DNA abundance levels ranged from 0.014 pg/ng to 25.7 pg/ng (1.4 x 10^-5^ to 2.57 x 10^-2^) in all inoculated seedlings. In contrast, no *C. weirii* was detected in any fine-root samples of non-inoculated control seedlings.

Regardless of the inoculation methods by either block-stick or dowel-plug, qPCR analysis of root collar samples revealed that all inoculated (both decayed and decay-free) trees were *C. weirii*-positive at 18 months post inoculation in the Yr-2017 trial. This indicates that the Yr-2017 trial had 100% infection rate, with visual decay rate at 40% and latent infection rate at 60%. *C. weirii* DNA abundance levels ranged from 0.049 pg/ng to 35.7 pg/ng (4.9 x 10^-5^ to 3.57 x 10^-2^) in the root collar samples (Fig. S3-B). In contrast, ITS-NGS analysis and visual wood-decay assessment detected fungal positive rate at 60% and 40%, respectively.

Similar to results from detection by ITS-NGS, qPCR detected *C. weirii* in the root collar of one non-inoculated control (Supplementary Fig. S3-B). There was no significant difference in fungal abundance between decayed and decay-free seedlings as detected by qPCR (Fig. 3B, t-test, p =0.28).

### Correlation of *C. weirii* relative abundances as detected by ITS-NGS and qPCR

The highest detection sensitivity for ITS-NGS was determined to be at 0.01% of targeted fungal reads relative to the total NGS reads while qPCR’s highest sensitivity was at 0.001% of targeted fungal DNA relative to the total DNA amount (ng/ng), with about 10 times increased sensitivity. Compared to *C. weirii-*detection methods by ITS-NGS and wood-decay symptom assessment, qPCR was demonstrated to have the highest detection level with the lowest false-negative rates. Following normalization of relative fungal abundance values by -log 10 (values + 10^-6^), *C. weirii* relative abundance values were highly comparable between detection methods of qPCR and ITS-NGS, with strong correlation (Pearson coefficient R = 0.86, p < 0.0001) (Fig. 4).

**Figure 4.**
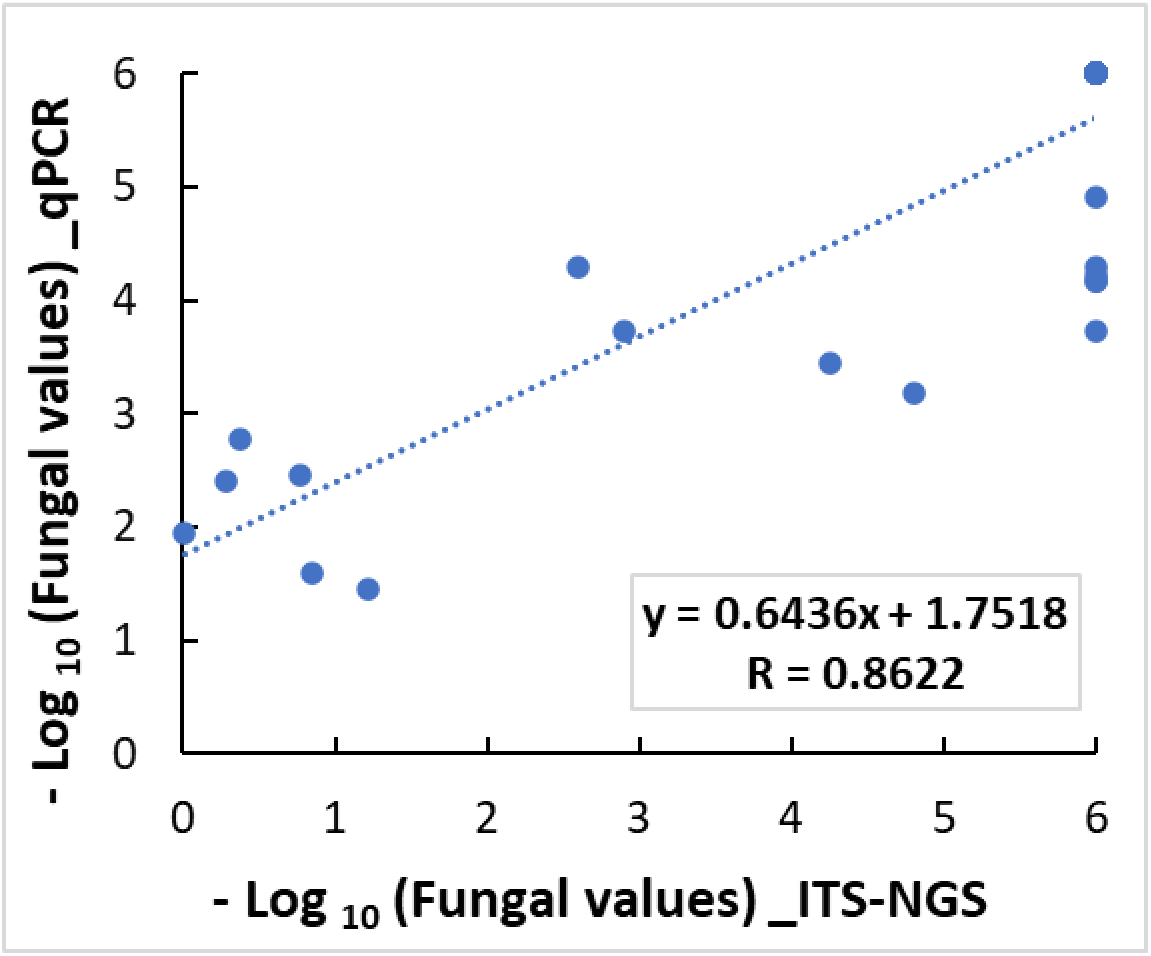
Comparison of *C. weirii* relative abundance as detected by qPCR and ITS-NGS from a total of 40 western redcedar tissue samples. Relative fungal abundance values were normalized to – log _10_ (values +10^-6^) before plotting Pearson’s correlation coefficient.

### Impacts of fungal inoculation on tree growth

Tree growth measurements revealed that inoculated seedlings had tree heights approximately 7%-10% less, and stem diameters approximately 6%-14% lower than the controls at 18-months post inoculation by all four fungi (Fig 5-A, 5-B). The inoculation impacts on tree height growth were significant for each of the fungi tested in this study (t-test, p < 0.05), but tree growth differences were not significant between the four tested fungi except that *H. occidentale*-inoculated seedlings had stem diameters smaller than either *C. weirii*- or *P. subacida*-inoculated seedlings (Fig. 5-B; t-test, p < 0.05).

**Figure 5.**
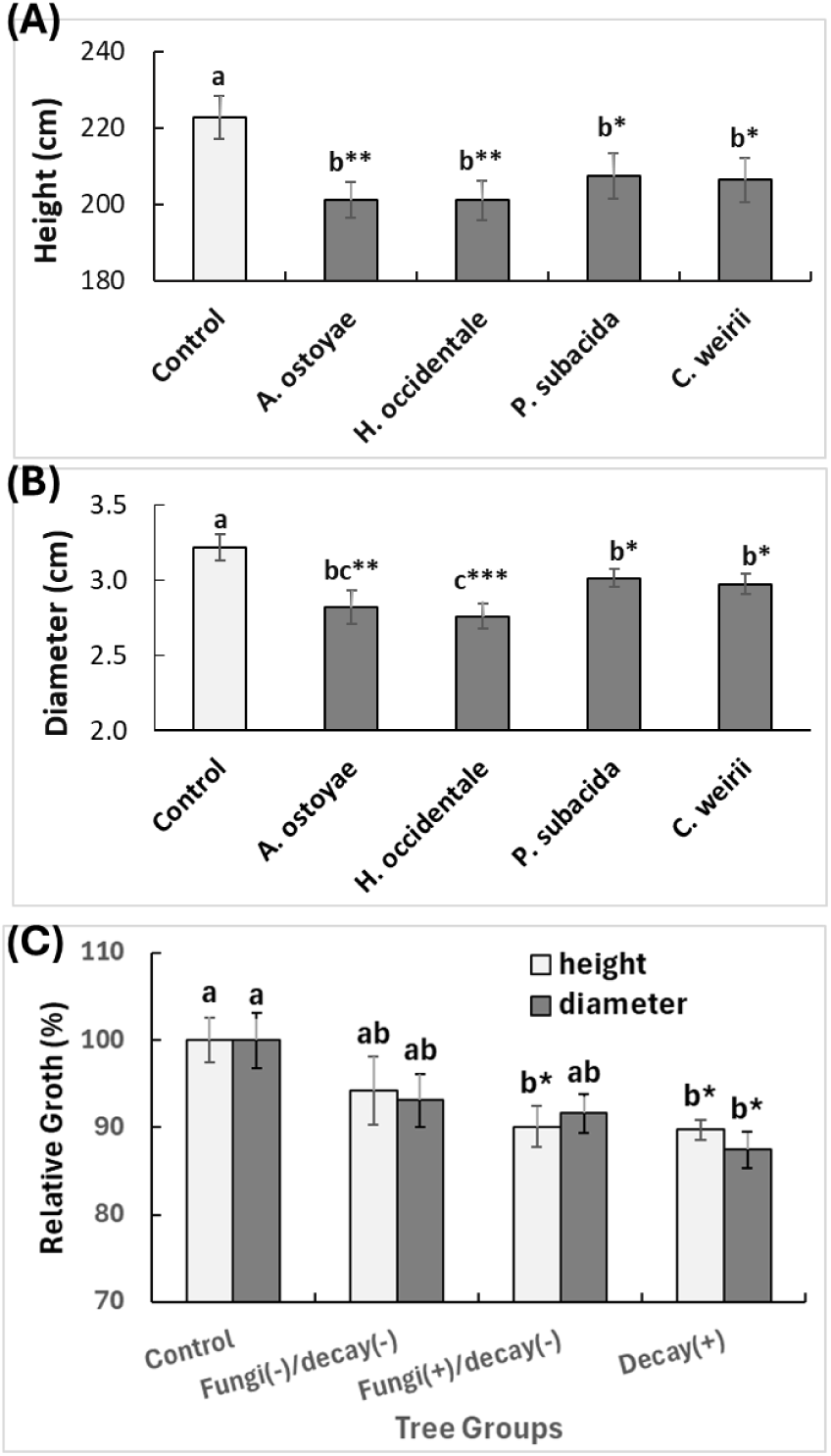
Tree growth inhibited by inoculation using root- and butt-rot fungi. (A) Seedling height and (B) stem diameters at ground level were measured at 18-months post inoculation. Vertical bars show the average values with standard error of mean (SEM). In (A) and (B), T-test was used to assess differences in seedling growth between the control and each decay-causing fungus separately. (C) Comparison of relative growths among tree groups using one-way ANOVA. Control represents uninoculated seedlings; Inoculated seedlings are presented in three groups. Fungi (-) and Fungi (+) represents absence or presence of targeted decay fungi as detected by ITS-NGS and qPCR; Decay (-) and Decay (+) represents absence or presence of visual wood decay. Stars indicate significant difference between treatments and controls with multiple comparison by the Bonferroni and Holm method (* p < 0.05).

Comparison of the two inoculation methods showed that inoculation by the dowel-plug method had a greater impact on seedling height than inoculation by the block-stick method, probably associated with wounding treatment (Fig. S1B). As compared to their controls, the heights and diameters decreased on average about 9.5% (t-test, p <0.001) and 11.7% (t-test, p < 0.01) respectively in dowel-plug-inoculated seedlings, while heights and diameters decreased on average about 4.6% (t-test, p = 0.177) and 11.6% (t-test, p < 0.05) respectively in block-stick- inoculated seedlings (Fig. S1B).

Based on the presence of wood decay and the presence of individual fungi in WRC tissues as detected by ITS-NGS and qPCR, infection rates were 100%, 90%, 80%, and 80% at 18 months post inoculation by *C. weirii*, *P. subacida*, *H. occidentale*, and *A. ostoyae*, respectively (Supplementary Table S4). Only five seedlings were not infected, two from inoculation using the block-stick method and three from inoculation using the dowel-plug method. All inoculated seedlings of Yr-2017 trial were divided into three groups: decayed, decay-free with latent infection, and absence of both decay and fungus although inoculated with no sign of infection. One-way ANOVA with post-hoc Tukey HSD Test revealed that decayed seedlings grew significantly slower (p < 0.05) in both heights (10% less) and stem diameters (13% less) than the non-inoculated controls. Although they were decay-free, those trees with latent infection had average height 10% shorter (p < 0.05), and stem diameters were 8% smaller than the non-inoculated controls but statistically not significant (p = 0.12). For the group of inoculated but not infected seedlings, their growth showed no significant difference from either the controls or pathogen-positive trees (Fig. 5-C).

## Discussion

### Controlled inoculation methods for effective assessment of WRC susceptibility and decay progression in response to decay fungi

The objectives for this study were to evaluate inoculation methods for different wood decay pathogens under controlled greenhouse conditions, and to investigate fungal detection methods for WRC. The block or block-stick method mimicked the natural transfer of the pathogen infection process via root contacts. The block and block-stick methods expose the host root cortical tissues to the fungus, while the dowel method bypass this. Further, the block- stick, or without stick method allows fungal soil and root colonization. The block placed in the top soil reduced the setup time considerably. With that in mind, successful fungal infection was confirmed by molecular detection of fungi in the WRC tissue samples for the four distinct fungal pathogens as well as by visual assessment of wood decay. Infection rates across all decay pathogens ranged from 80% to 100% but depended on the pathogen. The incidence in inoculated trees was comparable between the block (with or without stick) inoculation method and the dowel inoculation method. The incidence of visual wood decay varied largely from 20% to 60%, suggesting that the WRC seed families and seedlings may have variable susceptibility to different decay pathogens.

All four decay fungi were able to colonize healthy fine roots of WRC young seedlings using the block or block-stick inoculation method. Root to root contact is considered as the main pathway for root and butt rots to spread from infected roots to healthy living trees within a stand (Lewis 2013). This direct contact pathway is even more common for *C. weirii*, as its fruiting body is very rare in the field; and the fungal mycelia can grow along the WRC roots at yearly spread of 20–40 cm (Sinclair and Lyon 2005). Using the block inoculation method, it was found that *C. weirii* had a 30% early infection rate without development of decay at 6-months post inoculation. Furthermore, at 18-months post inoculation, the root infection rate increased to 100%, and the subsequent decay rate reached 17%, similar to a previous report (Sturrock and Reynolds 1998). The fungal basidiospores may spread through wind and water to stems or healthy tree roots, while wound sites are possible entry points for spores (Hagle 2009). This could explain why one uninoculated seedling was infected by *C. weirii* in our greenhouse Yr- 2017 trial.

Although WRC infection rates were at high levels up to 100% and visual decay rates at the root collar were at intermediate levels up to 60%, none of the infected seedlings were killed by any of the four tested wood decay fungi, and no external disease symptoms were found. In contrast to another tree species, western hemlock (*Tsuga heterophylla*) seedlings inoculated with *H. occidentale* showed high root and butt rot incidence rates with 7% mortality and 43% expressing clear disease symptoms (such as needle-shed, chlorotic foliage, reduction in leader and branch growth) at 8 months post controlled inoculation by a wound-based method (Liu et al. 2022). Previous field surveys (Theis and Sturrock 1995) also found low WRC mortality rate as compared to other conifers in younger stands. In older stands where disturbance was rare, stem decay led to stem breakage and contributed significantly to mortality (Hennon 1995).

Evidence from these studies confirmed that WRC had high incidence but low mortality due to the wood decay pathogens. We detected rates of decay incidence up to 60% in root collars of five-year old WRC seedlings. Although no morphological disease symptoms were observed in our inoculation trial, all wood decay fungi had about 10% impact on seedling height growth, similar to basal area loss in another field study of older WRC trees (Morrison et al. 2014). Both seedlings inoculated with decay and seedlings that were infected but decay-free grew slower than non-inoculated seedlings, indicating that latent infection by decay fungi significantly affected growth of WRC seedlings.

### Advantages and limitations of ITS-NGS for diagnosis of WRC wood decay diseases

The second main objective of this study was to develop molecular tools for fast and reliable detection of decay pathogens in WRC living tissues. We demonstrated an integration of ITS-NGS and qPCR for the study of fungal infection biology and profiling of fungal communities in the WRC tissues. For molecular diagnostics, this study first estimated the accuracy and utility of ITS-NGS using the Illumina MiSeq platform for detection of the targeted decay fungi. NGS of the ITS regions is considered as the best option to analyze fungal communities with wide applications, and so far, no viable alternative to ITS has been well developed for broad community analyses of fungi (Schoch et al. 2012). In contrast to qPCR for measurement of absolute abundance of known pathogens, NGS analysis of ITS regions through (meta-) barcoding in a large set of samples has the potential to provide information on relative abundance of individual OTU in the complete community of organisms, including all culturable and non-culturable known and unknown species (Müller et al. 2023). Of two ITS regions (ITS-1 and ITS-2), this study selected ITS-1 as target for NGS because ITS-1 has better distinction among species and better performance in terms of richness and taxonomic coverage (Nilsson et al. 2008; Mbareche et al. 2020). ITS-NGS was a powerful detection method for targeted pathogens in both decayed and decay-free WRC seedlings, but with lower detection rates of fungal presence in all decayed wood. This limitation using decayed wood samples may be caused by biases of ITS-NGS that are commonly linked to some taxonomic groups due to primer mismatch and variations of copy number, AT/GC content, and length of ITS regions (Bellemain et al. 2010; Kauserud 2023). In addition, WRC tissue types and gDNA extraction methods likely affected efficiency and accuracy of microbial profiles generated by ITS-NGS. For example, DNA extraction from WRC decayed wood was more difficult than that from fine roots. Methods of DNA extraction favoured some microbial species and types of tissues which influenced microbial community profiles (Frau et al. 2019). In addition to sequencing ITS-1, further NGS analysis of ITS-2 or long-read sequencing of the full length of the ITS region may help with the detection of more wood decay pathogens and the diversity of the fungal community present in WRC samples. Overall, despite limitations and potential biases in ITS amplification using universal fungal primers, NGS of fungal ITS regions has provided a global estimation of fungal incidence in WRC by different decay pathogens as well as an overview of the microbiome associated with roots and decayed wood tissues.

Counts of each unique ITS seq from NGS data are not suitable for quantification of absolute levels of different fungal species within sampled tissues (Shelton et al. 2022; Kauserud 2023). However, our data showed that the normalized counts of ITS sequences can be used for quantitative comparisons of the relative abundance of the same fungal species across different samples. As analysed using *C. weirii* as the target, this ITS-NGS-based relative fungal abundance was highly correlated with that by qPCR measurement, consistent with a recent report (Shelton et al. 2022). Integration of ITS-NGS with in-vitro fungal culture and qPCR analyses is suggested for examination of the role of rDNA dynamics in fungal community dynamics (Lavrinienko et al. 2021). This genomics approach adapted for investigation of WRC decay fungi may be applicable to the study of infection biology and the microbiota in other forest pathosystems and ecosystems, allowing future exploration of the relationship between root microbiota and forest health, particularly in field studies.

### Development of species-specific qPCR assays for fast and reliable diagnosis of *C. weirii* in WRC samples

We developed the first set of qPCR diagnostic assays through amplification of six genomic regions to enable detection and quantification of *C. weirii,* the most important white rot stem decay pathogen in WRC stands. The novel *C. weirii*-specific qPCR assays developed in this study were aimed at providing fast and reliable molecular diagnostic tools for monitoring infection of WRC trees. As suggested by the results of the fungal ITS-NGS analysis, a relevant number of important and widespread wood decay fungi are detectable using the ITS-NGS approach. However, this NGS-based diagnostic approach requires professional staff with costly instruments and bioinformatic pipeline for data analysis, hence, it may not be appropriate for most plant disease diagnostic laboratories. To its advantage, the qPCR assays are better to quantify abundance of the fungal targets for better evaluation of the proportion of targeted pathogens inside the host WRC tissues, without a concern for potential biases associated with the ITS-NGS-based detection method as noted above.

This study standardized the qPCR assays for quantitative detection of *C. weirii* in WRC tissues and discriminated it from *C. sulphurascens* and other wood-decay pathogens. Although *C. weirii* may have evolutionarily separated from *C. sulphurascens* about 38 Mya with considerable genomic divergency (Leal et al. 2019), few molecular diagnosis tools have been reported for species identification in the genus *Coniferiporia*. One set of ITS primers was previously developed for endpoint PCR to distinguish *C. weirii* from *C. sulphurascens* (Lim et al. 2005). More recently, an ITS-based multiplex PCR assay was also available to detect the *C. weirii/sulphurascens* complex from other groups of decay fungi commonly found in conifers, but can not distinguish C. weirii from C. sulphurascens (Gonthier et al., 2015). Both *C. weirii* and *C. sulphurascens* are pathogens of conifer trees causing laminated root rot in North America and eastern Asian countries (Lewis 2013), but *C. weirii* mainly infects WRC while *C. sulphurascens* mainly infects Douglas fir (*Pseudotsuga menziesii*) in North America (Larsen et al. 1994). As compared to endpoint PCR, qPCR provides greater insight into community function and dynamics of microbiomes associated with plant pathosystems. However, due to high conservation of related sequences, ITS sequence divergence is usually not sufficient to discriminate between the closely related taxa. Furthermore, large intraspecific variation in rDNA copy number complicates analysis of rDNA amplicon data, and it is still largely unknown how eukaryote microbial communities alter their rDNA copy number in response to environmental changes (Lavrinienko et al. 2021). These facts limit the ITS region as a suitable genomic target of qPCR assays of individual fungal species.

In this study, we searched other genomic regions for proper species-specific variations to design species-specific PCR primers through sequence alignment analysis among related fungal species. The qPCR primer sets developed in this study were first tested for their *C. weirii*- specificity by endpoint PCR using pure gDNA extracted from cultured mycelia of related root and butt rot fungi. Their diagnostic efficiency was further validated through generation of standard curves using pure fungal gDNA. These qPCR assays demonstrated a high degree of repeatability for detection and quantification of *C. weirii* associated with WRC tissues.

### qPCR assays for monitoring infection pathways and phytosanitary purposes

Of six qPCR assays developed in this study, the qPCR assay of Cw-RPBb (F76/R235) showed amplification efficiencies > 90%, which is in the efficiency range for properly developed PCR assays (Roche 2000). Regardless of inoculation methods and WRC tissue types, the qPCR assay of Cw-RPBb detected *C. weirii* infection at frequencies much higher than ITS-NGS. Using the same set of WRC tissue samples, the qPCR assay demonstrated about 10 times higher sensitivity than ITS-NGS. Compared to ITS-NGS, qPCR assays have an advantage of absolute quantification of pathogens inside the host tissues with lower cost, simple operation, and without biases associated with the NGS-based detection method.

One picogram of DNA is 978 megabases long and the *C. weirii* genome is about 42.2 megabases long (McMurtrey et al. 2023). The lowest detectable gDNA amount in a qPCR reaction is 0.2 pg, which approximately equates 4-5 copies of the *C. weirii* genome. Evidence from these analyses indicates that this qPCR assay enables the sensitive, specific, and accurate quantification of viable cells as low as 2∼3 *C. weirii* cells inside WRC infected tissues. Of all tested wood-decay samples, *C. weirii* gDNA was reliably detected by the qPCR assay as low as 1.4 x 10^-5^ of total DNA in nanograms. The qPCR assays developed here showed high detection sensitivity, close to the theoretical maximum limit and comparable to the best ones previously reported for other pathogens.

Because of their high specificity, reliability, and sensitivity, our qPCR assays can be used to detect trace amounts of *C. weirii* in field samples for disease surveillance and diagnosis. Long distance spread of decay pathogens is a serious concern when wood, bark, and seedlings are transported. *C. weirii is* recommended for quarantine by the European and Mediterranean Plant Protection Organization (EPPO 2014). Traditional in-vitro fungal culture techniques are limited in detecting trace amounts of living fungus in infected samples, and this traditional approach is time-consuming causing delays in response. Alternatively, the qPCR assay helps to identify the infection pathway of *C. weirii* in soils and other environmental and timber samples more rapidly for phytosanitary purposes to reduce this delay in response. At a subsequent stage, this qPCR assay can be further developed into a portable tool for diagnostic utility in field surveys.

### Species-specific qPCR assays detected early and latent infection with potential application for breeding WRC quantitative disease resistance

Currently, WRC susceptibility to biotic agents is still assessed through conventional inoculation trials or field tests (Russell and Yanchuk 2012). Our controlled inoculation trials showed that WRC seedlings developed wood decay diseases at 18-months post successful infection by four soil-born decay pathogens. Even under greenhouse conditions, resistance screening through assessment of wood-decay is time-consuming, requiring at least 5 years of seedling growth and intensive labor work for greenhouse maintenance. Evaluation of early traits with the help of marker-assisted selection (MAS) still needs the development of molecular tools for screening young seedlings before resistant genotypes can be selected, characterized, and utilized in existing breeding programs.

Latent infection phase has been observed in several tree species post infection by various pathogens (Negahban et al. 2024). Presence of a few fungal cells in host tissues (such as fine roots or barks) may indicate the fungal penetration sites allowing early, initial infection before decay development. These initial infected pathogens can be latent in host tissues for a long period of time with multiple years prior to development of disease symptoms under favorable environmental conditions, such as abiotic drought and temperature stresses (Luo et al. 2024). Field surveys suggest that the wood decay-causing fungi can survive in a latent phase within WRC tissues for extended periods before causing wood decay and killing trees (Morrison et al. 2001; Konchalski 2015; Cruickshank et al. 2018). Recently a qPCR method was reported to detect early infection (Chandelier et al. 2019), as well as latent infection of pathogens in asymptomatic trees of several wood species (Luo et al. 2024; Negahban et al. 2024). Our *C. weirii* block inoculation trials allowed for fungal movement through the soil to infect roots, in some of which the qPCR assay was able to detect infection without visual wood decay symptoms at a 30% and 60% incidence rate in trees at 6- and 18-months post inoculation, respectively. Longer latent period is one of the most important polygenic traits involving a diversity of different mechanisms for host quantitative disease resistance against various pathogens (Bove and Rossi 2020). For a given host-pathogen complex in a controlled environment, a longer latency period for disease symptoms indicates a higher level of host disease resistance. As compared to decayed seedlings, those infected but decay-free seedlings likely had latent infection, which might indicate disease resistance, or that they might not express decay symptoms at all during their lifespan. Slowing decay progression upwards through the stem heartwood would reduce the amount of cull at harvest.

A limited or slower spread of pathogens from root infection sites up to trunk wood tissues may indicate quantitative disease resistance that has restricted movement of the invaded pathogen. Wood tissues are commonly drilled out from trunks or cut down from smaller branches for molecular detection of wood pathogens (Avenot et al. 2022). Gradient of fungal levels from root collar to upper positions of the trunk can be tested using qPCR as biomarkers to determine the speed of pathogen spread, which may be used as an index of WRC susceptibility to predict development of decay. Our qPCR assays have been validated for effective detection of early or latent infection of *C. weirii* in both fine roots and stem wood tissues of asymptomatic WRC seedlings. MAS tools like these qPCR assays can accelerate resistance screening of WRC at an early stage of selection. A future study will be performed to determine the correlation between fungal DNA levels in host tissues at early or latent infection stages and the final disease symptom of wood-decay.

In conclusion, artificial inoculation methods used in this study were effective for four decay fungi to infect the WRC young seedlings. These provide practical protocols for screening WRC genotypes with enhanced resistance to these and other related root and butt rot diseases in breeding programs. We recommend the block method over the dowel stem method for pathological studies because the infection occurs by root penetration through the cortical tissues, similar to natural infection. High throughput ITS-NGS and qPCR assays were developed with highly comparable results for the qualitative and quantitative characterization of wood decay fungi in infected WRC living tissues. We highlighted the potential of ITS-NGS and qPCR as integrative molecular approaches for further studies of infection biology of WRC wood-decay diseases. The newly developed qPCR assays can then be more effective with lower cost for screening for disease resistance by tree breeders, and detection of targeted pathogens in environmental samples and natural stands. Monitoring decay and disease infection processes will enhance our understanding of fungal infection courts and mechanisms, and host resistance against root and butt rot disease among cedar populations.

## Acknowledgements

We thank Rona Sturrock and Kevin Pellow for the setup of inoculation trial. We thank the BC Ministry of Forests for seedlings and seeds used in the study, and their long-term tree breeding program for making this possible. We thank the CFS-PRM program for funding awarded to JJL, and to the Pacific Forestry Centre for providing greenhouse space.

## Author contributions

J.-J.L and M.C. designed the research. S.H. and A.Z. performed the experiments. J.-J.L, M.C. and A.Z. collected greenhouse data. J.-J.L. and S.H. analyzed data and prepared the manuscript. A.Z., I.L., and C.F. revised the manuscript. All authors contributed to the article and approved the submitted version.

---

**Figure S1.**
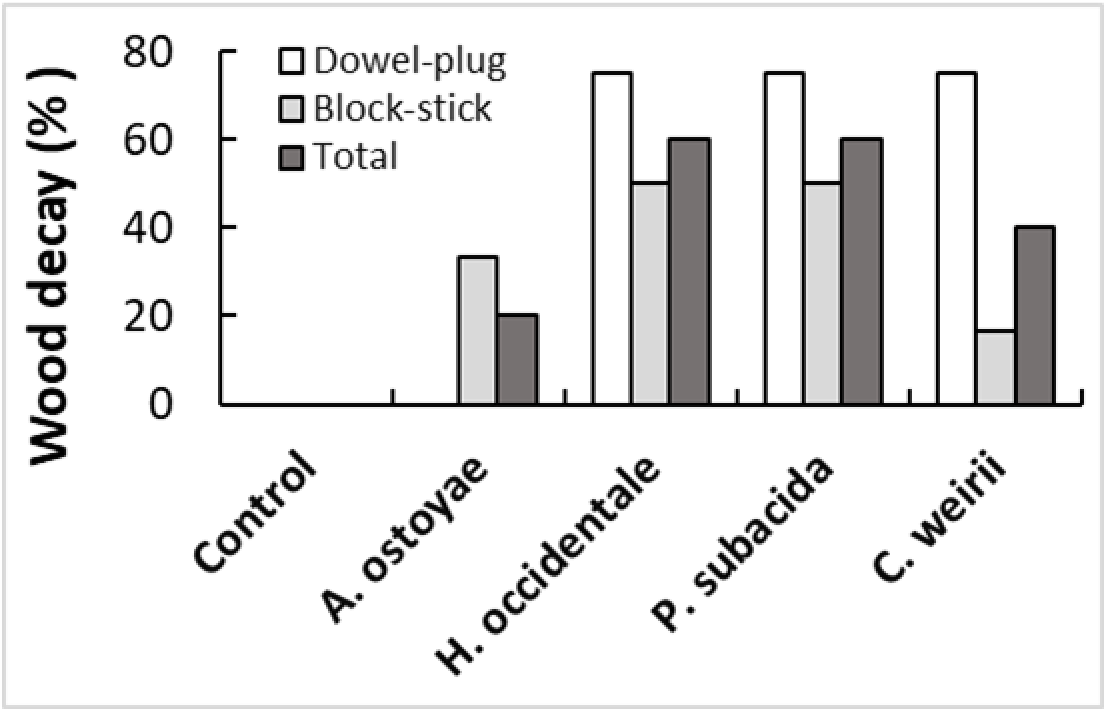
Comparison of two inoculation methods used in the Yr-2017 trial. (A) Visual wood decay incidence of western redcedar trees inoculated with four fungi. The control and the dowel method for *A. ostoyae* had zero decay. (B) Seedling growth impacted by fungal inoculations. Seedling height and stem diameter were individually measured at 18-months post inoculation. Vertical bars show relative growth in averages with standard error of mean (SEM). Relative growth of seedlings inoculated using block-stick and dowel-plug methods were compared with their corresponding controls (Control-1 and -2), and significant differences were tested using the t-test (* p < 0.05, ** p < 0.01, *** p <0.001).

**Figure S2.**
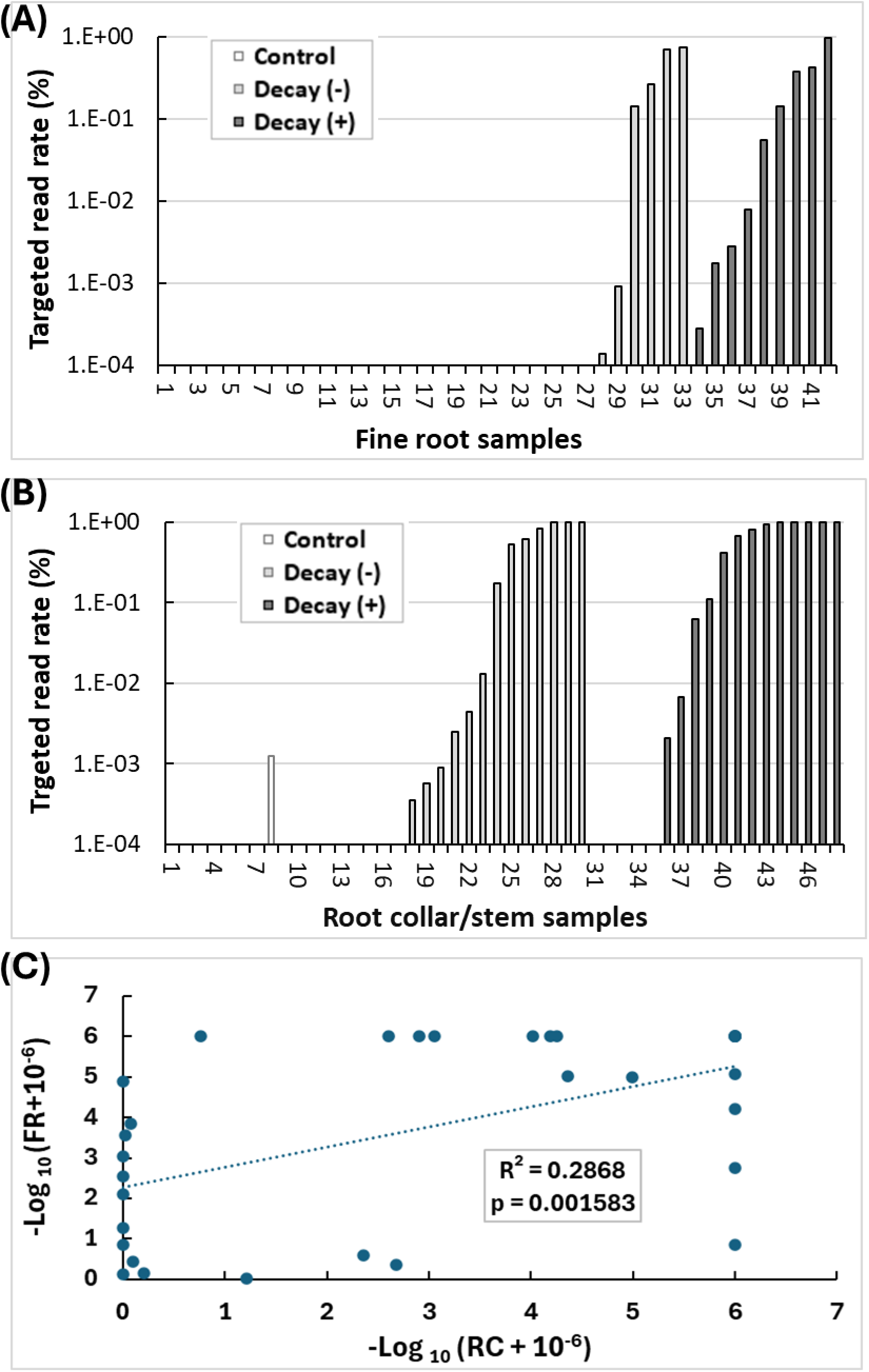
Relative abundance of targeted fungi as detected by ITS-NGS. Percentages of targeted fungal reads in total ITS-NGS reads were calculated for each tissue samples of western redcedar seedlings. (A) Fine root samples from 42 seedlings. Sample Nos. 1-8 are uninoculated seedlings (controls) with pathogen-positive rate 0%; samples 9-33 are inoculated but decay-free seedlings (Decay -) with pathogen-positive rate 24%; samples 34-42: decayed seedlings (Decay +) with pathogen-positive rate 100%. (B) Root collar samples from 48 seedlings. Samples No.1-8 are uninoculated seedlings with pathogen-positive rate 12.5%; samples 9-30 are inoculated but decay-free seedlings with pathogen-positive rate 59.1%; samples 31-48 are decayed seedlings with pathogen-positive rate 72.2%. (C) Correlation of relative fungal abundances between root collar (RC) and fine root (FR) samples of the same set of seedlings. Relative fungal abundance values were normalized to - log10 (values +10^-6^) before calculating Pearson’s correlation coefficient.

**Figure S3.**
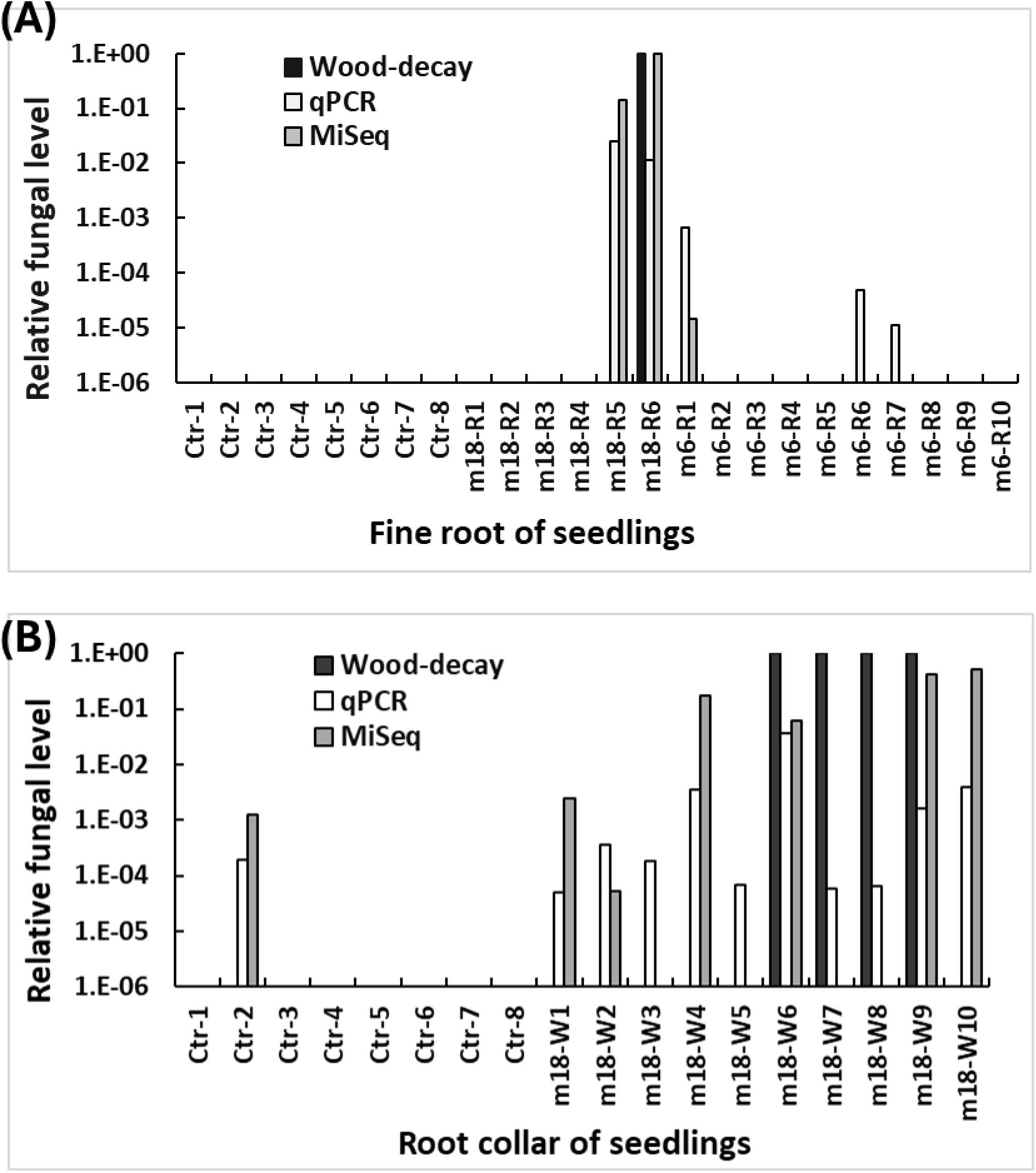
Infection of western redcedar tissues by *C. weirii* as detected by ITS-NGS, qPCR, and visual wood-decay. (A) Fine root samples of 24 seedlings, including 8 non-inoculated as controls (Ctr-1 to Ctr-8), 6 inoculated seedlings samples at 18-months post inoculation (m18-R1 to m18-R6), and 10 seedlings at 6 months post inoculation (m6-R1 to m6-R10). (B) Root collar samples of 18 seedlings at 18 months, including 8 non-inoculated as controls (Ctr-1 to Ctr-8) and 10 inoculated seedlings (m18-W1 to m18-W10) using visual symptoms, PCR or ITS-NGS.

**Figure S4.**
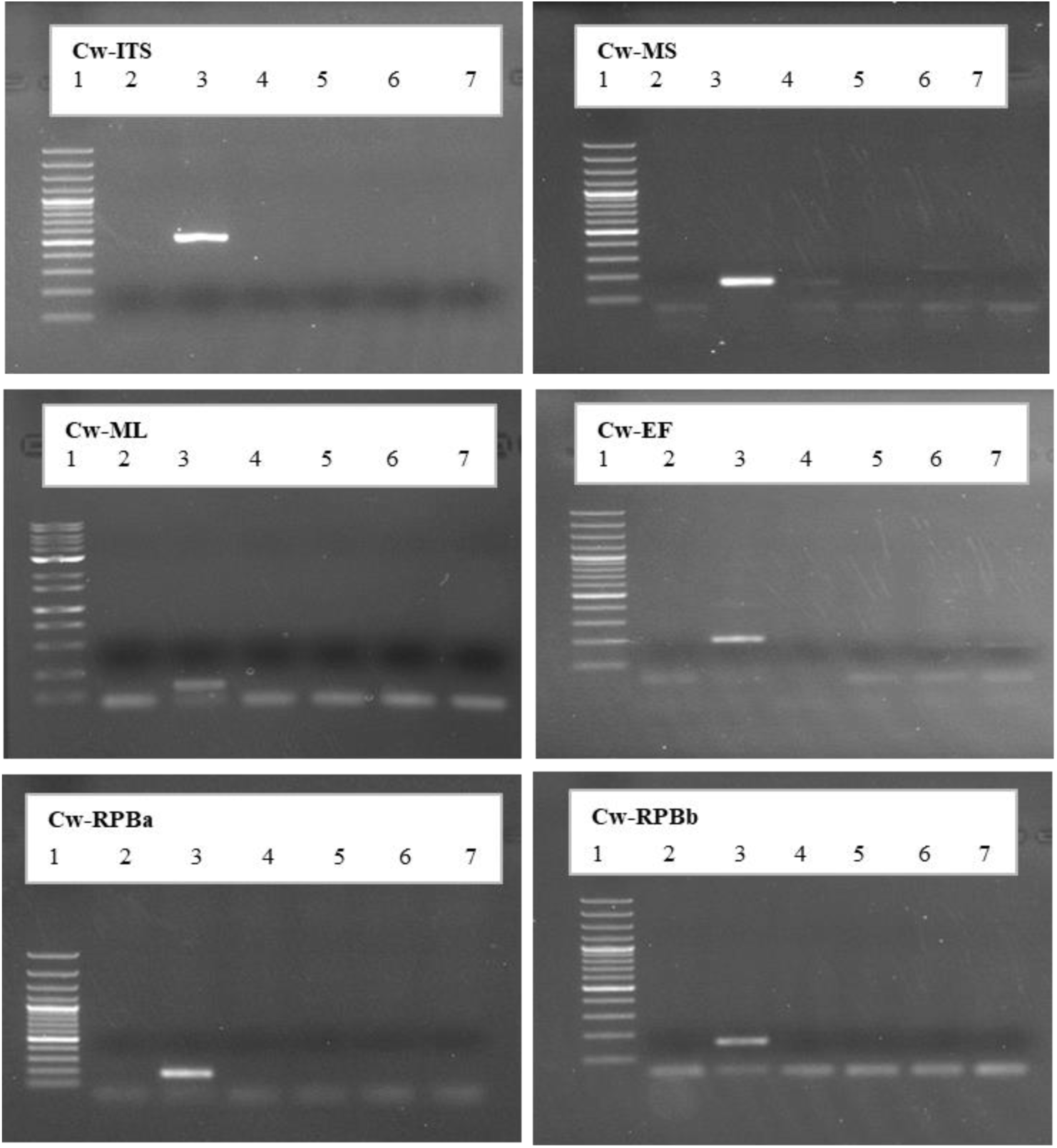
Agarose gel electrophoresis showing the amplicons of endpoint PCR reactions. Six genomic regions were amplified using PCR primers designed for *C. weirii.* Genomic DNA was extracted from different decay fungi as PCR templates. Lane 1: Fermentas GeneRuler 100bp plus DNA ladder, Lane 2: negative control; Lane 3: *C. weirii*, Lane 4: *C. sulphurascens*, Lane 5: *P. subacida;* Lane 6: *H. occidentale*, Lane 7: *A. ostoyae*.

Figure S5. Amplification profiles (left) and standard curve analysis (right) of qPCR assays for detection of *C. weirii*. Right panels show the linear relationship between the log10 of known amount of *C. weirii* gDNA in tenfold dilutions and Ct value of qPCR amplicon. The qPCR assay was run in technical triplicate using SYBR Green Master Mix with the taxon-specific primer.

Table S1. Sample sizes of western redcedar seedlings consisting of two full sibling families and two seedlots, two inoculation techniques, and four decay pathogens.

Table S2. Decay fungi used in this study.

Table S3. Nucleotide sequences of PCR primers for molecular diagnosis of decay fungi.

Table S4. Successful infection and disease incidence rates.

Table S5. Specificity of *C. weirii* qPCR assays as tested by endpoint PCR.

Table S6. Ct values of *C. weirii* qPCR assays and melting temperatures (Tm) of their amplicons.

